# Probing the sub-cellular mechanisms of LCA5-Leber Congenital Amaurosis and associated gene therapy with expansion microscopy

**DOI:** 10.1101/2023.01.17.524360

**Authors:** Siebren Faber, Olivier Mercey, Katrin Junger, Alejandro Garanto, Marius Ueffing, Rob W.J. Collin, Karsten Boldt, Paul Guichard, Virginie Hamel, Ronald Roepman

**Affiliations:** Department of Human Genetics, Radboud Institute for Molecular Life Sciences, Radboud University Medical Center, Nijmegen, the Netherlands; Department of Molecular and Cellular Biology, University of Geneva, Geneva, Switzerland; Division of Experimental Ophthalmology and Medical Proteome Center, Center of Ophthalmology, University of Tübingen, Germany; Department of Pediatrics, Radboud Institute for Molecular Life Sciences, Radboud University Medical Center, Nijmegen, the Netherlands; Department of Human Genetics, Donders Institute for Brain, Cognition and Behaviour, Radboud University Medical Center, Nijmegen, the Netherlands

## Abstract

Leber Congenital Amaurosis (LCA) is a group of Inherited Retinal Diseases (IRDs) characterized by the early onset and rapid loss of photoreceptor cells. Despite the discovery of a growing number of genes associated with this disease, the molecular mechanisms of photoreceptor cell degeneration of most LCA subtypes remain poorly understood. Here, using retina-specific affinity proteomics combined with Ultrastructure Expansion Microscopy (U-ExM), we revealed the structural and molecular defects underlying LCA type 5 (LCA5) with unprecedented resolution. We showed that *LCA5*-encoded lebercilin, together with Retinitis Pigmentosa 1 protein (RP1) and the intraflagellar transport (IFT) proteins IFT81 and IFT88, localize at the bulge region of the photoreceptor outer segment (OS), a region crucial for OS membrane disc formation. Next, we demonstrated that mutant mice deficient for lebercilin exhibit early axonemal defects at the bulge region and the distal OS, accompanied by reduced level of RP1 and IFT proteins, affecting membrane disc formation and presumably leading to photoreceptor death. Finally, we probed the *LCA5* gene augmentation therapy strategy using U-ExM by monitoring its subcellular outcome. We found that, expression of *LCA5* partially restores the bulge region, preserves OS axoneme structure and membrane disc formation, as well as photoreceptor survival.

## Introduction

Mutations in the *LCA5* gene disrupt its cognate ciliary protein lebercilin (herein refer to as LCA5 for simplicity), causing Leber congenital amaurosis (LCA), one of the most severe forms of inherited blindness (1). Phenotypically, LCA manifests as nystagmus, delayed pupillary responses, photophobia, hyperopia, and severely decreased visual acuity, already in the first year of life (2). The cells that are primarily affected in LCA5 patients are the light-sensing photoreceptors, the most abundant cell type in the human retina (1, 3). Photoreceptors are highly specialized neuroepithelial cells, consisting of structurally and physiologically distinct cellular compartments, including a synaptic terminal, an inner segment (IS), an outer segment (OS), and a connecting cilium (CC) that bridges the IS and OS (4).

LCA5 has been shown to be widely expressed in human and mouse tissues, notably at the level of ciliated epithelia (1, 5, 6). However, despite this ubiquitous pattern of expression, *LCA5* mutations lead to a retina-restricted phenotype in human, mouse and zebrafish (1, 7, 8), suggesting a crucial function of this protein in the retina. In the mouse eye, *Lca5* mRNA is increasingly expressed in photoreceptor cells during development (1), arguing for a specific role of LCA5 in these cells. At the subcellular level, LCA5 has been described as a microtubule-binding protein, which localizes at the base of the primary cilium in epithelial cells (1, 7). In line with this, LCA5 interacts with several centriole/basal body and ciliary components such as OFD1 (9), NINL (10) or intraflagellar transport (IFT) core proteins (7). Based on regular immunohistochemistry, LCA5 has been proposed to localize to the photoreceptor CC, a highly developed equivalent of the ciliary transition zone involved in selective transport of OS proteins, lipids, and membrane vesicles (1, 7, 11, 12). Recently, we optimized super-resolution ultrastructure expansion microscopy (U-ExM) for retinal imaging to get new insights into the molecular architecture of photoreceptor OS with unprecedented resolution. This technique, based on the isotropic physical expansion of a biological sample embedded in a swellable gel, revealed that LCA5 localizes to a specific photoreceptor region, that we called ‘bulge region’, apical to the CC (13), in accordance with our previous EM analysis (1). This region exhibits a stereotypical enlargement of the axonemal microtubules compared to the CC, correlating with the absence of the inner scaffold, a recently described molecular zipper maintaining cohesion between microtubule doublets (MTDs) at the level of the CC and the centrioles (13, 14). Currently, there is not much known about the exact function of the bulge region. It is suggested that it is important for the initiation of OS membrane disc formation in an actin-dependent manner, mediated by PCARE and WASF3 (15). Given the retina-specific phenotype observed with mutations of *LCA5*, unraveling the function of the bulge protein LCA5, could bring insights into the role of the bulge region in context of specific forms of inherited retinal diseases (IRDs).

In addition to deciphering the molecular mechanisms associated with IRDs, one of the most exciting challenges in recent years in the ophthalmologic field is the development of therapeutic strategies to restore vision in the context of these diseases, that are to date mostly incurable. Gene augmentation therapy has emerged as an attractive tool for monogenic IRDs, because the eye is easily accessible for delivery injection techniques, immune privileged, and highly suitable for assessing functional outcomes (16-20). Commonly, delivery of a gene therapy is performed by intravitreal or subretinal injection of a functional copy of the affected gene, encapsulated by a viral vector (21). Until now, Luxturna is the only FDA-approved gene therapy amenable for LCA patients with biallelic *RPE65* mutations, although several other gene therapies are currently in clinical trials (21, 22). Before getting to the clinical phase, pre-clinical animal studies are essential for the development of safe and effective gene therapies (21). The quality of the outcome measures of these animal studies are therefore of utmost importance to enhance the success rate in later stages. The methods that are used to assess the therapeutic efficacy of the gene therapy in most pre-clinical gene therapy studies are mainly focused on the assessment on tissue level and visual acuity, including electroretinography, optical coherence tomography, and retinal immunostainings (23-26). Although these methods are valuable tools to assess gene therapy efficacy in mice, we are currently lacking assays to evaluate the efficacy of gene augmentation therapy at the photoreceptor cellular and ultrastructural level. Here, we propose a novel and easily accessible approach based on U-ExM to tackle this question at a nanoscale resolution, using a previously described gene therapy for LCA5 (23, 24) as proof-of-concept.

## Results

### Affinity proteomics of LCA5 in mouse retina reveals photoreceptor-associated modules

To better understand the molecular mechanisms leading to Leber congenital amaurosis, we first investigated the retina-specific interactome of LCA5 in both *Lca5*^gt/gt^ (HOM) and *Lca5*^*+/gt*^ (HET) mice. Hence, we used an AAV7m8 vector to deliver a human *LCA5* cDNA construct (23, 24), encoding LCA5 fused at the C-terminus with a 3xFLAG-tag, into the eyes of the mice at P2 by intravitreal injection. Four weeks post-injection, anti-Flag affinity purification was performed followed by mass spectrometry analysis.

Analysis of the LCA5-associated proteins in injected HOM mice revealed a total of 108 significantly enriched proteins (Figure 1A and Supplemental Table 1). Analysis of the LCA5-associated proteins in injected HET mice showed a highly similar set of significant proteins (Supplemental Figure 1 and Supplemental Table 1), confirming the robustness of our approach.

**Figure 1.**
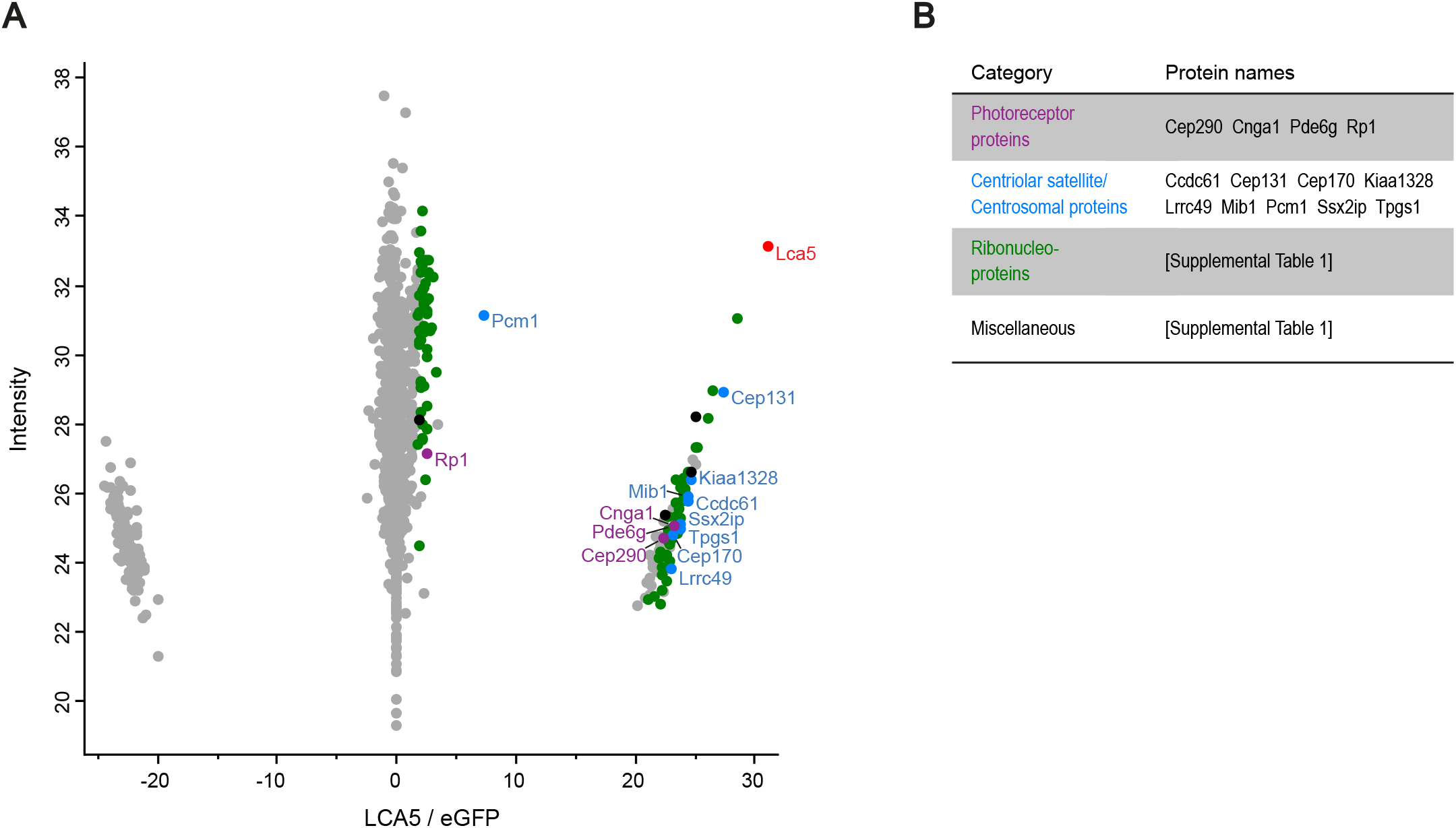
Identification and clustering of potential LCA5 interactors. (**A**) Scatterplot showing enriched proteins, comparing AAV-*LCA5* injected retinas to AAV-*eGFP* (control) injected retinas in *Lca5gt/gt* (HOM) mice. The bait protein (LCA5) is shown in red. Significantly enriched proteins are categorized into different groups based on their function, including photoreceptor-associated proteins (purple), centriolar satellite/centrosomal proteins (blue), ribonucleoproteins (green), and miscellaneous (black). X-axis represents log2 ratio between AAV-*LCA5* and AAV-*eGFP* (control) injected retinas. Y-axis represents the intensity score, indicating the relative amounts of proteins in the dataset. (**B**) Table showing the significant proteins categorized in different groups, based on their function. Original data is listed in Supplemental Table 1.

The identified significant proteins can be categorized into different groups based on their functions. These groups include: photoreceptor-associated proteins, centriolar satellite/centrosomal proteins, ribonucleoproteins, and miscellaneous (Figure 1B, Supplemental Figure 1B, and Supplemental Table 1).

Since LCA5 is a ciliary protein with a prominent role in photoreceptors (1, 7), we focused on the photoreceptor-associated proteins and the centriolar satellite/centrosomal proteins identified in our datasets (Figure 1, A and B). More specifically, we further investigated the central player in the centriolar satellite/centrosomal proteins group, PCM1 (27, 28), and two photoreceptor-associated proteins, CEP290 and RP1. Interestingly, we recently found that LCA5 localizes in the extension of CEP290 (13), suggesting an interdependence between these proteins, and mutations in *CEP290* lead also to Leber congenital amaurosis (29). Similarly, RP1 has been shown to localize at the distal part of the connecting cilium (CC) (30, 31), similar to LCA5, and mutations in *RP1* lead to retinitis pigmentosa (32, 33).

### LCA5 is a bulge protein localizing in between CEP290 and RP1

To precisely map these potential LCA5 interactors in photoreceptor cells at high resolution, we emphasized on super-resolution U-ExM, which we recently optimized for expanding mouse retinal tissue (13). Consistent with our previous work (13), we found LCA5 predominantly at the proximal part of the bulge region, with an additional weaker signal detected at the distal axoneme and the centrioles (Figure 2A). Since LCA5 shows faint localization at the centrioles, we first investigated the photoreceptor-specific localization of the centriolar satellite protein PCM1, which is also involved in the correct localization of several proteins to the centrioles (34). As expected, we found that PCM1 localizes to the centriolar satellites, but we could not detect it at the bulge region (Supplemental Figure 2A), suggesting that LCA5 and PCM1 might interact at the level of satellites rather than along the photoreceptor axoneme. To address this point, we co-stained PCM1 and LCA5 in human U2OS cultured cells (Supplemental Figure 2B), and found that both co-localize at the level of centriolar satellites.

**Figure 2.**
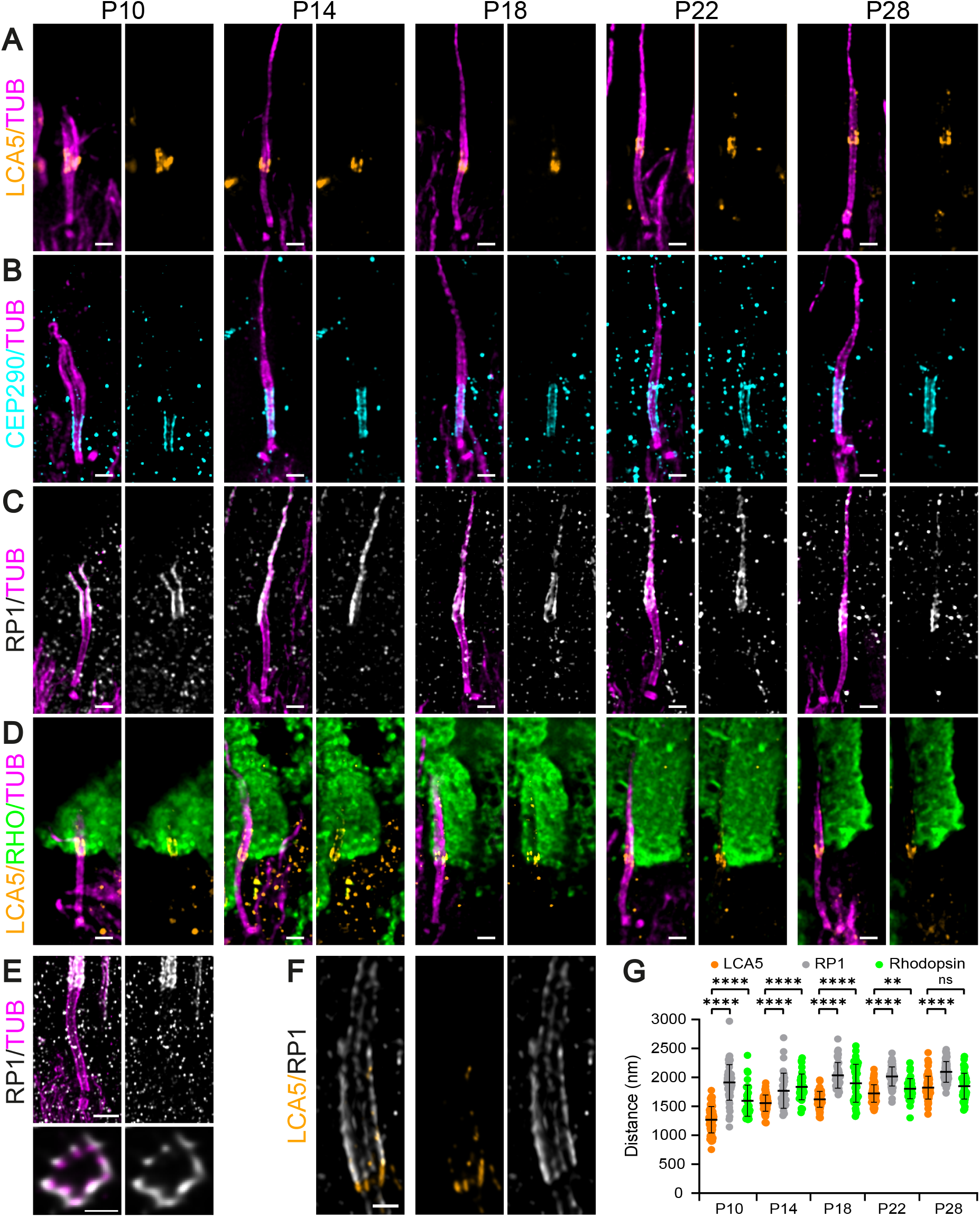
Nanoscale mapping of LCA5. (**A-D**) Widefield (63x) images of expanded photoreceptors stained for tubulin (magenta) and LCA5 (orange, A), CEP290 (cyan, B), RP1 (white/gray, C), or LCA5/Rhodopsin (orange/green, D) from P10 to P28 in *Lca5*^*+/gt*^ (HET) mice. Scale bars: 500 nm. (**E**) Confocal U-ExM images of adult photoreceptor stained for tubulin (magenta) and RP1 (white/gray). Lower panels show transversal view of the bulge region. Scale bars: 500 nm (side view); 200 nm (transversal view). (**F**) Confocal U-ExM image of adult photoreceptor stained for LCA5 (orange) and RP1 (white/gray). Scale bar: 500 nm. (**G**) Quantification of the distance of LCA5 (orange), RP1 (white/gray), and Rhodopsin (green) signal proximal ends to the mother centriole proximal end from P10 to P28. Three animals per timepoint.

Next, we set out to investigate the specific location of CEP290, and RP1 in the developing photoreceptors from P10 until P28 in *Lca5* HET mice. CEP290 was consistently detected at the distal end of the daughter centriole and the external part of the microtubules along the CC (Figure 2B), confirming that LCA5 localizes in the extension of CEP290 (13). Measuring the lengths of CEP290 and the inner scaffold protein POC5 along the CC showed that their signal gradually increases during photoreceptor development, reaching a plateau phase from P18 until P28, with an average final length of 1420 nm (CEP290) and 1552 nm (POC5), similar to what we previously reported (13) (Figure 2B and Supplemental Figure 3B).

**Figure 3.**
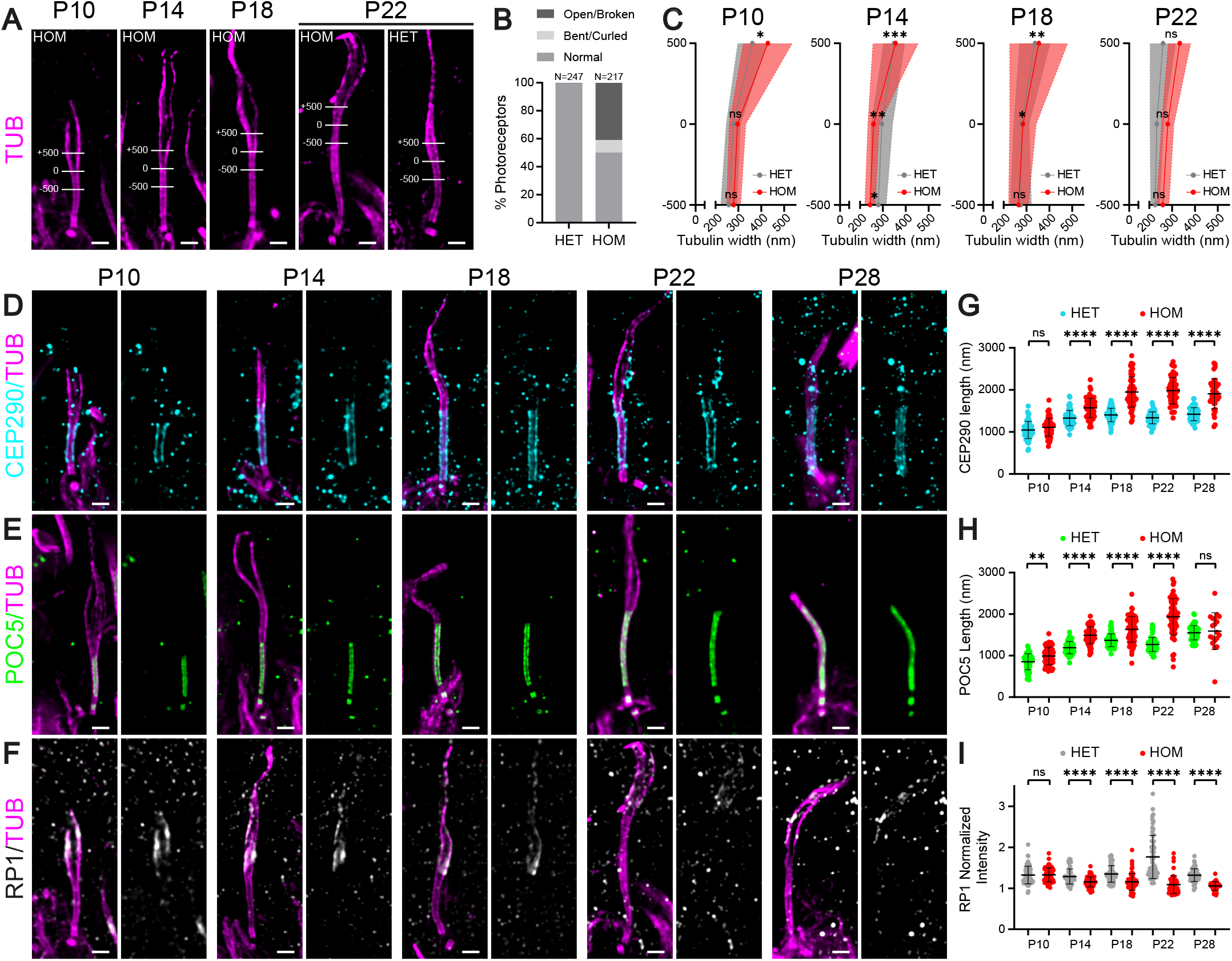
Effect of LCA5 loss on bulge formation, distal axoneme organization and CC length. (**A**) Widefield (63x) images of expanded photoreceptors stained for tubulin (magenta) from P10 to P22 in *Lca5*^*gt/gt*^ (HOM) mice and in P22 *Lca5*^*+/gt*^ (HET) mice. Lines in P10 image illustrate the measurements used in (C) of tubulin width at 3 locations. 0: the distal end of CC inner scaffold marker POC5 was used to set the 0 location; −500: 500 nm proximally to the CC inner scaffold end; +500: 500 nm distally to the CC inner scaffold end. Scale bars: 500 nm. (**B**) Distal axoneme (above CC) conformations of HET vs. HOM photoreceptors at P18 and P22 indicated in percentages. Photoreceptors are categorized in three different distal axoneme conformations, including Normal (HET: 99.6%; HOM: 50.2%), Open/Broken (HET: 0.4%; HOM: 40.8%), and Bent/Curled (HET: 0%; HOM: 9%) axonemes. Number of photoreceptors analyzed: N=247 (HET); N=217 (HOM). (**C**) Tubulin width measurements of the photoreceptor at the 3 locations depicted in (A) (−500 nm, 0 nm, +500 nm) from P10 to P22. Average tubulin width at each location (− 500 nm, 0 nm, +500 nm) is indicated by a gray or red dot for HET and HOM, respectively. Measurements for P28 were not included since most of the distal axonemes are lost at this timepoint. Only photoreceptors that were stained for tubulin and POC5 were used for the measurements. Three animals per timepoint. Asterisks indicate level of significance for the tubulin width dispersion between HET and HOM (not the difference in average tubulin width between HET and HOM). (**D-F**) Widefield (63x) images of expanded photoreceptors stained for tubulin (magenta) and CEP290 (cyan, D), POC5 (green, E), and RP1 (white/gray, F) from P10 to P28 in HOM mice. Scale bars: 500 nm. (**G-I**) Impact of LCA5 loss on CEP290 length (G), CC inner scaffold length (POC5, H), or RP1 normalized intensity at the bulge region (I) from P10 to P28. Three animals per timepoint.

Interestingly, we found that RP1 localizes to the distal part of the bulge region, where it displays a stereotypical 9-fold symmetry on the external part of the MTDs similarly to LCA5 (13), and also extends towards the distal axoneme (Figure 2, C and E). Moreover, co-stainings of LCA5 and RP1 revealed that RP1 is continuous to the LCA5 signal (Figure 2F). We next mapped its position relative to LCA5 by measuring the distance from the centriolar distal end to the proximal end of the RP1 and LCA5 signals. We demonstrated that the RP1 signal is significantly higher compared to LCA5 at all timepoints (Figure 2G). Remarkably, the RP1 signal distance is significantly longer at P10 compared to P14 (Figure 2G; p=0.0005), possibly explained by the immaturity of the axoneme at P10, including the bulge region. Previously, it was shown that RP1 regulates distal axoneme formation and is required for proper OS formation (30, 35). Therefore, we investigated the localization of the initiation site of the OS by staining the OS marker rhodopsin relative to LCA5. We found that LCA5 localizes next to the proximal extremity of the rhodopsin staining (Figure 2D). By measuring the proximal end distance of the rhodopsin signal, we found that it coincides at almost all timepoints with the region in between the proximal signals of LCA5 and RP1 (Figure 2G), suggesting that the bulge region might indeed be crucial for OS membrane disc formation.

### LCA5 loss affects CC length, bulge formation, and distal axoneme organization

To better understand the role of the bulge region in photoreceptor development, and more specifically the function of LCA5 at this location, we made use of an *Lca5*^*gt/gt*^ (HOM) mouse model (7). Phenotypically, this mouse model fully recapitulates LCA in human, showing an early onset, progressive retinal degeneration, with only a few layers of photoreceptor nuclei left in the outer nuclear layer (ONL) at P28 (7). It is noteworthy that assessment of the P28 timepoint in HOM retinas is rather difficult, since most photoreceptors, and especially their distal axonemes, are already lost at this timepoint.

First, we looked at the impact on photoreceptor integrity by staining the axonemal microtubules using U-ExM in HET versus HOM mice. We found that approximately 50% of photoreceptors of HOM mice show distal axoneme abnormalities with a loss of cohesion of the microtubules above the CC, including open/broken and bent/curled conformations (Figure 3, A and B). To validate the observed loss of cohesion in the distal axoneme, we measured the axoneme diameter at different positions: at the distal end of the CC (0), 500 nm below in the CC (−500) and 500 nm above the CC (+500) in both HET and HOM mice (Figure 3, A and C). We found a significant increase in axoneme diameter above the CC (+500 nm) in HOM mice at P10 (p=0.0468) and P22 (p<0.0001), but not at P14 (p=0.6842) and P18 (p=0.7434). However, we did find a significant bigger dispersion of tubulin width in the HOM photoreceptors at P10, P14 and P18 compared to HET mice (Figure 3C), suggesting that loss of LCA5 either leads to the collapse or the spreading of microtubules above the CC.

Based on the tubulin staining, the CC seems unaffected by the loss of LCA5. To confirm this, we stained for the CC proteins CEP290 and POC5, and found that they are properly localized in absence of LCA5 (Figure 3, D and E). However, when measuring the lengths of the CEP290 or POC5 signal along the CC, we found them to be significantly increased from P14 to P28 (Figure 3, G and H), indicating that the overall CC length increases in absence of LCA5 and of the bulge region.

Since the distal axoneme is clearly disrupted by the loss of LCA5, we hypothesized that RP1 will also be affected. Indeed, we could show that in absence of LCA5, RP1 levels at the bulge region, while not yet affected at P10, are significantly decreased at almost every timepoint, with almost no signal left at P28 (Figure 3, F and I).

### LCA5 loss affects rhodopsin localization

We showed that LCA5 localizes next to the proximal part of the rhodopsin staining and that loss of LCA5 leads to the disruption of the distal axoneme, the microtubule-based cytoskeleton of the OS. To further investigate the impact of LCA5 loss on OS development, we examined the OS protein rhodopsin over time. We first analyzed rhodopsin and tubulin at low magnification (Supplemental Figure 4). We found a significant decrease in ONL thickness over time, with only a few layers of photoreceptor nuclei left at P28 (Supplemental Figure 4, B and C). Moreover, rhodopsin was mis-localized to the ONL (Supplemental Figure 4, B and D), confirming our previous observations (7). To get a better understanding of the rhodopsin mis-localization, we looked at rhodopsin and tubulin at high magnification (Figure 4, A and B). In contrast to CEP290 and RP1, which appear unchanged at P10, we observed that rhodopsin localization starts to be affected at P10, with a clear accumulation of rhodopsin above the basal body (Figure 4B, white arrows), suggesting a defective rhodopsin trafficking along the photoreceptor axoneme. At this timepoint, the rhodopsin staining at the OS level does not seem to be impaired, but this rapidly exacerbates from P14 onwards (Figure 4B). Besides the accumulation of rhodopsin above the basal body, we observed rhodopsin along the CC (Figure 4B, white arrowheads) and in membrane vesicle-like structures (Figure 4B, open white arrowheads), further supporting the rhodopsin transport defect hypothesis, which precedes the CC over-extension and the distal axoneme disruption.

**Figure 4.**
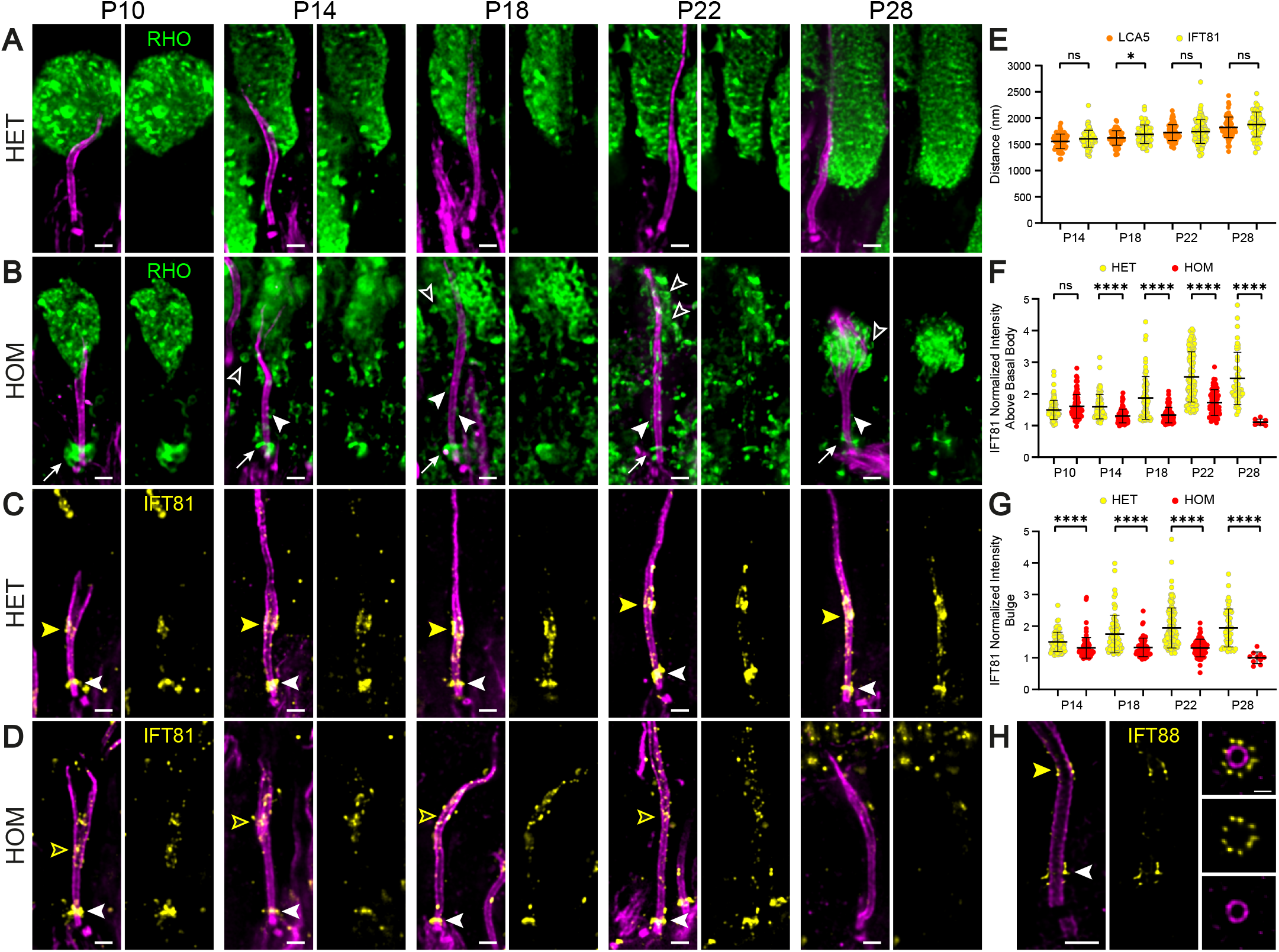
Effect of LCA5 loss on OS formation and intraflagellar transport. (**A-B**) Widefield (63x) images of expanded photoreceptors stained for tubulin (magenta) and Rhodopsin (green) from P10 to P28 in *Lca5*^*+/gt*^ (HET) (A) and *Lca5*^*gt/gt*^ (HOM) (B) mice. White arrows indicate accumulation of Rhodopsin above the basal body. White arrowheads indicate Rhodopsin along the CC. Open white arrowheads indicate Rhodopsin in vesicle-like structures. Scale bars: 500 nm. (**C-D**) Widefield (63x) images of expanded photoreceptors stained for tubulin (magenta) and IFT81 (yellow) from P10 to P28 in HET (C) and HOM (D) mice. White arrowheads indicate IFT81 localization above the basal body. Closed and open yellow arrowheads indicate IFT81 localization at the bulge region in HET and HOM, respectively. Scale bars: 500 nm. (**E**) Quantification of the distance of LCA5 (orange; same values as in Figure 2G) and IFT81 bulge region (yellow) signal proximal ends to the mother centriole proximal end from P14 to P28. Measurements for P10 were not included since the bulge region is not properly formed yet at this timepoint. Three animals per timepoint. (**F-G**) Impact of LCA5 loss on IFT81 normalized intensity above the basal body (F) or at the bulge region (G) from P10 to P28. Measurements for P10 were not included for IFT81 intensity at the bulge region since it is not properly formed yet at this timepoint. Three animals per timepoint. (**H**) Confocal U-ExM images of adult photoreceptor stained for tubulin (magenta) and IFT88 (yellow). Right panels show transversal view of the bulge region. White arrowhead indicates IFT88 localization above the basal body. Yellow arrowhead indicates IFT88 localization at the bulge region. Scale bars: 500 nm (side view); 200 nm (transversal view).

### LCA5 regulates IFT-B trafficking at the bulge region

To investigate the potential trafficking defect, we next investigated the precise localization of the IFT-B protein IFT81 in photoreceptors of HET mice, followed by analyzing the alterations seen in photoreceptors of HOM mice over time. In HET mice, IFT81 accumulates above the basal body and at the bulge region, but also to a lesser extend along the CC (Figure 4C). To assess whether IFT81 and LCA5 localize to the same proximal region at the bulge, we measured the distance from the centriolar proximal end to the proximal end of both signals and found that IFT81 and LCA5 localize to the same position in the bulge with the exception of the P18 timepoint (Figure 4E). Moreover, we found that IFT88, another IFT-B protein displays a similar localization at the bulge, but also localizes at the base of the cilium with a 9-fold symmetry, similarly to what has recently been described in the green algae *Chlamydomonas* (36) (Figure 4H).

Next, we analyzed the effect of LCA5 loss on IFT81 localization. We observed a significant decrease in IFT81 levels above the basal body from P14 onwards to P28 (Figure 4, D and F). IFT81 localization was similarly decreased at the bulge at every timepoint, and we noticed that the signal is dispersed along the distal axoneme before vanishing at P28 (Figure 4, D and G).

Taken together, our results demonstrate that the loss of LCA5 affects the localization of IFT81, which we postulate to result in the mis-trafficking of rhodopsin leading to defective OS membrane disc formation and ultimately photoreceptor cell degeneration.

### Probing LCA5 gene augmentation therapy with U-ExM reveals partial rescue of photoreceptor cells

Capitalizing on the molecular and structural understanding of the LCA5-associated disease gained using U-ExM, we next decided to probe the AAV-*LCA5* gene augmentation therapy (23, 24) to further improve its efficacy.

Since LCA5 has been found to be expressed 15 days after intra-ocular delivery (23), we focused our analysis on the P18, P22, and P28 timepoints. Intriguingly, when dissecting the retinas, we noticed that the retina did not fully recover its normal aspect, as seen by the degree of pigmentation, suggesting that the gene augmentation was probably only partial (Supplemental Figure 5A).

**Figure 5.**
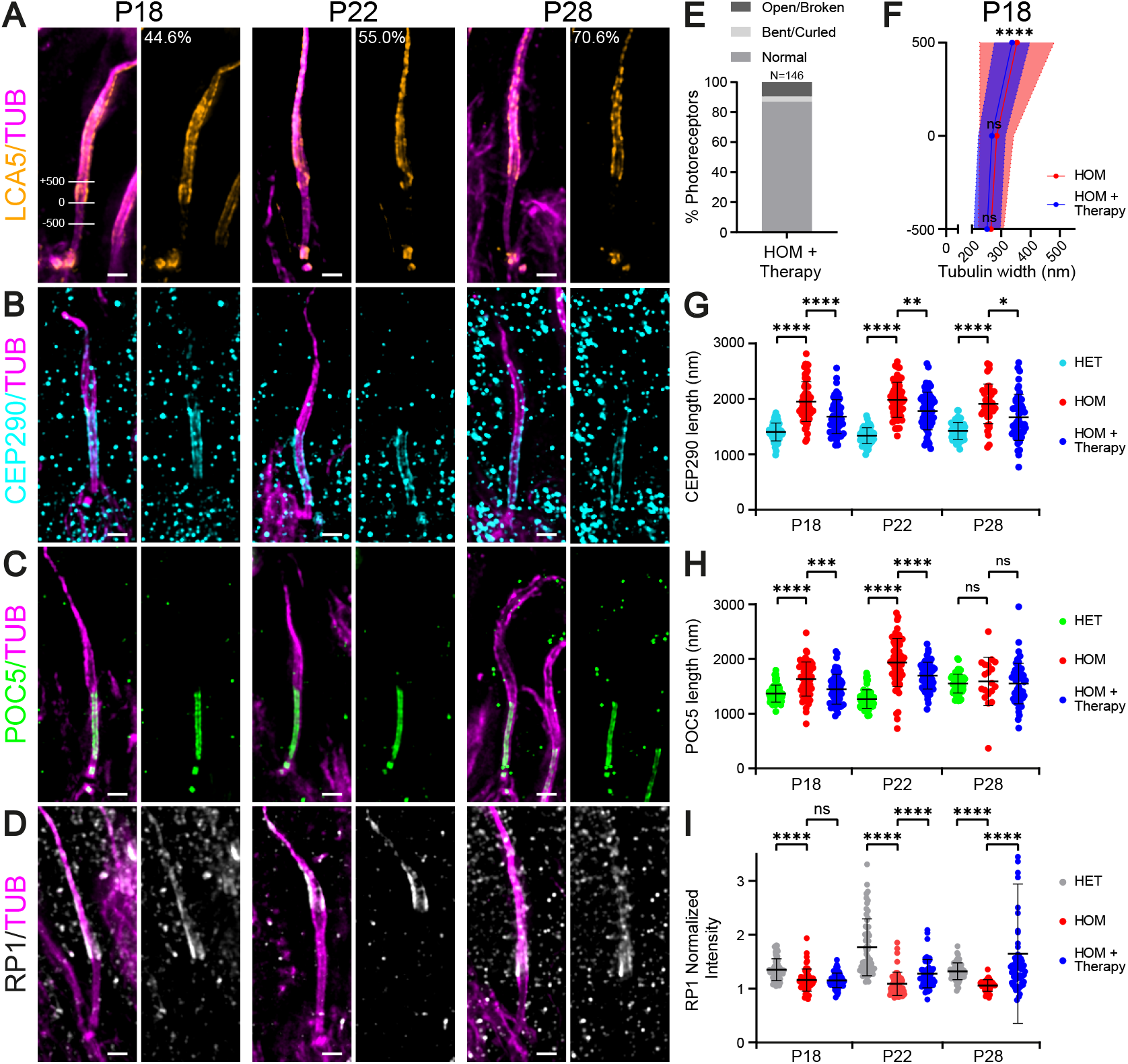
Effect of AAV-*LCA5* gene augmentation therapy on distal axoneme organization and CC length. (**A-D**) Widefield (63x) images of expanded photoreceptors stained for tubulin (magenta) and LCA5 (orange, A), CEP290 (cyan, B), POC5 (green, C), or RP1 (white/gray, D) from P18 to P28 in AAV-*LCA5* gene therapy treated *Lca5*^*gt/gt*^ (HOM) mice. Lines in P10 LCA5 image (A) illustrate the measurements shown in (E) of tubulin width at 3 locations. 0: the proximal end of the LCA5 signal was used to set the 0 location; −500: 500 nm proximally to the LCA5 proximal end; +500: 500 nm distally to the LCA5 proximal end. Percentages shown in (A) indicate the percentage of photoreceptors that re-express LCA5 at each timepoint. Three animals per timepoint. Scale bars: 500 nm. (**E**) Distal axoneme (above CC) conformations of AAV-*LCA5* gene therapy treated HOM photoreceptors from P18 to P28 indicated in percentages. Photoreceptors are categorized in three different distal axoneme conformations, including Normal (87%), Open/Broken (9.6%), and Bent/Curled (3.4%) axonemes. Number of photoreceptors analyzed: N=146. (**F**) Tubulin width measurements of P18 HOM photoreceptors gene therapy treated vs. non-treated at the 3 locations depicted in (A) (−500 nm, 0 nm, +500 nm). Average tubulin width at each location (−500 nm, 0 nm, +500 nm) is indicated by a red or blue dot for HOM and HOM + therapy, respectively. Measurements for P28 were not included, since most of the distal axonemes are lost in HOM mice at this timepoint. Only photoreceptors that re-express LCA5 were used for the measurements. Note that HOM measurements correspond to the data presented in Figure 3C. Three animals per timepoint. Asterisks indicate level of significance for the tubulin width dispersion between HOM and HOM + therapy (not the difference in average tubulin width between HOM and HOM + therapy). (**G-I**) Impact of AAV-*LCA5* gene therapy on CEP290 length (G), CC inner scaffold length (POC5, H), or RP1 normalized intensity at the bulge region (I) from P18 to P28. Note that HET and HOM measurements correspond to the data in Figure 3, G-I. Three animals per timepoint.

Consistently, LCA5 was detected in 44.6%, 55.0%, and 70.6% of P18, P22, and P28 photoreceptors, respectively (Figure 5A). The gradual increase in percentage of LCA5-positive photoreceptors could possibly be explained by the increasing loss of photoreceptors that were not targeted by AAV-*LCA5* at later timepoints. At low magnification, ectopic LCA5 expression was observed through all retinal layers (Supplemental Figure 5B). In photoreceptors, LCA5 localization was restored, but not restricted to the bulge region, as it was found along the entire distal axoneme (Figure 5A). Furthermore, LCA5 seemed enriched at the centrioles compared to the endogenous expression of LCA5 (Figure 5A). These results demonstrate that LCA5 ectopic expression, while partially restoring the normal LCA5 localization, also induces non-expected distribution, which might impact the full functional rescue.

Next, we evaluated the effect of *LCA5* augmentation on axonemal photoreceptor axoneme rescue, combining all three timepoints (P18, P22 and P28). Importantly, we found that more than 87% of the LCA5-positive photoreceptors contained a normal-shaped distal axoneme (Figure 5E). Furthermore, looking at LCA5-positive photoreceptors, we found that the dispersion in tubulin width above the CC is significantly reduced at P18 in the gene therapy treated retinas (Figure 5F), indicative of a partial rescue of axonemal defects upon LCA5 expression.

Next, we monitored the effect of the gene augmentation therapy on the distribution of the CC proteins CEP290 and POC5 in the entire photoreceptor population (including the positive and negative LCA5 cells). We found that both CEP290 and POC5 signal lengths were significantly reduced in the AAV-*LCA5* treated retinas (Figure 5, B, C, G and H), but not fully restored to the normal length.

We next tested whether RP1 distribution is restored upon LCA5 expression. We found that RP1 localization was not restored at P18 (Figure 5, D and I). At P22 and P28 however, the RP1 signal at the distal axoneme was significantly rescued compared to the non-injected group (Figure 5, D and I), indicating that exogenous LCA5 expression is sufficient to drive proper RP1 localization.

Finally, we investigated whether *LCA5* augmentation had an effect on OS restoration and photoreceptor viability. To do so, we first measured the ONL thickness and found a modest but significant increased ONL thickness at P22 and P28 (Figure 6, A and D). Moreover, rhodopsin localization was partially restored at P18 and P28, with reduced mis-localization at the ONL level (Figure 6, A and E). Importantly, at high magnification, part of the photoreceptors exhibits total OS restoration, with rhodopsin localizing in a rod-shaped manner and no accumulation above the basal body at all timepoints (Figure 6B). Despite these promising results, we noticed large heterogeneity in OS restoration between replicates, possibly due to the variability the injection procedure, the number of targeted photoreceptors or the LCA5 expression levels (Supplemental Figure 5C).

**Figure 6.**
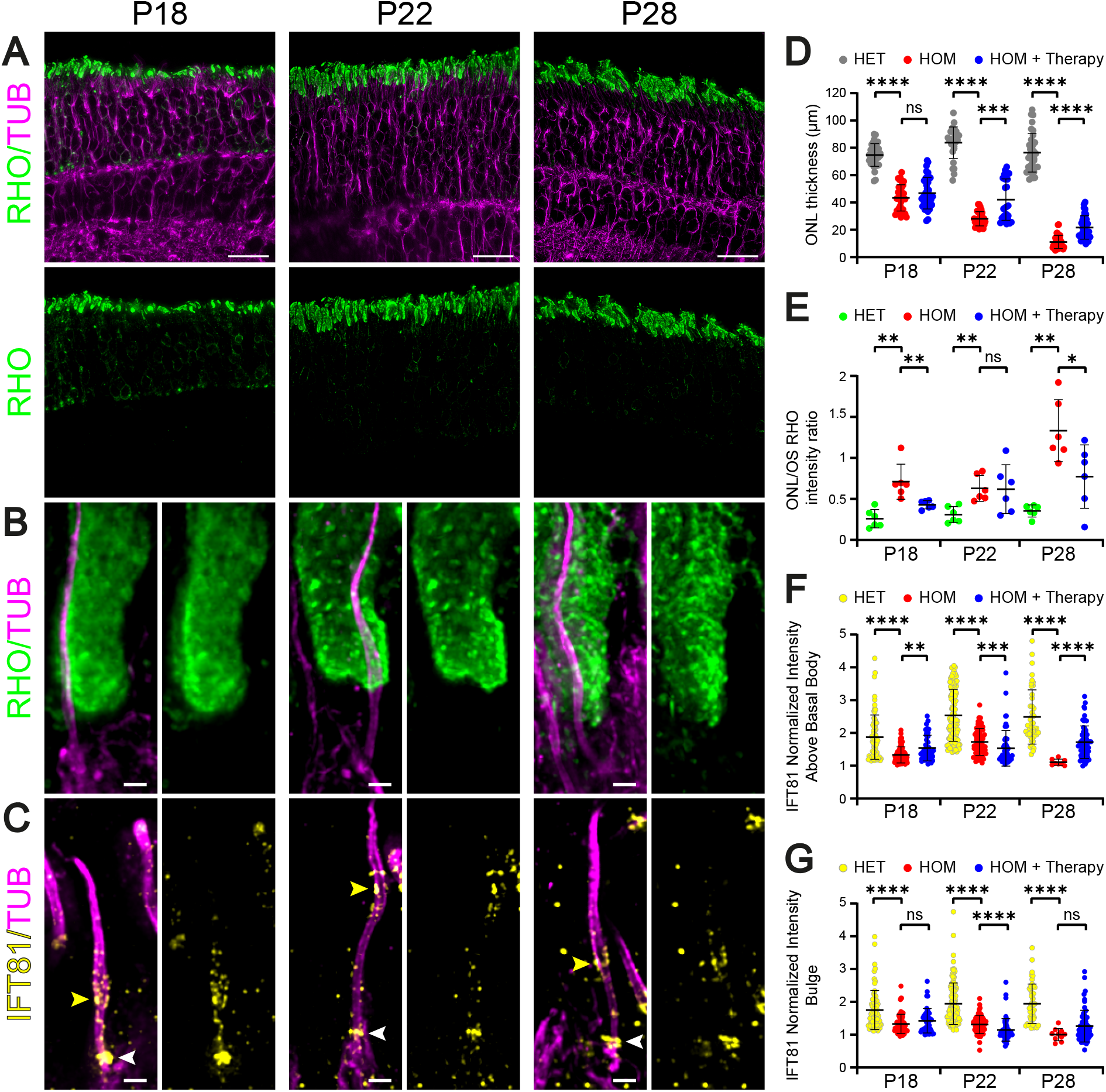
Effect of AAV-*LCA5* gene augmentation therapy on OS formation and intraflagellar transport. (**A**) Low magnification (20x) widefield images of expanded *Lca5*^*gt/gt*^ (HOM) retinas, treated with AAV-*LCA5* gene therapy, showing rod OS restoration from P18 to P28 by staining with Rhodopsin (green) and tubulin (magenta). Scale bars: 20 μm. (**B-C**) Widefield (63x) images of expanded photoreceptors stained for tubulin (magenta) and Rhodopsin (green, B) and IFT81 (yellow, C) from P18 to P28 in AAV-*LCA5* gene therapy treated HOM mice. White arrowheads in (C) indicate IFT81 localization above the basal body. Yellow arrowheads in (C) indicate IFT81 localization at the bulge region. Scale bars: 500 nm. (**D-G**) Impact of AAV-*LCA5* gene therapy on outer nuclear layer (ONL) thickness (D), ONL/OS Rhodopsin intensity ratio (E), IFT81 normalized intensity above the basal body (F), or IFT81 normalized intensity at the bulge region (G) from P18 to P28. Note that HET and HOM measurements in (F) and (G) correspond to data in Figure 4, F and G. Three animals per timepoint.

The above results suggests that rhodopsin trafficking is partially restored by gene therapy. To confirm that trafficking is restored, we monitored IFT81 localization at the bulge region and above the basal body. At the bulge region, IFT81 levels were slightly increased after gene therapy at P18 and P28, although not significant (Figure 6, C and G). In contrast, IFT81 levels at P22 were reduced by AAV-*LCA5* treatment compared to non-treated photoreceptors (Figure 6, F and G), probably due to the heterogeneity of LCA5 expression inside the retina and the ectopic expression along the distal axoneme (Figure 5A and Supplemental Figure 5A). Above the basal body, the levels of IFT81 were significantly increased by the LCA5 augmentation at P18 and P28 (Figure 6, C and F), suggesting a partial rescue of IFT81, primarily above the basal bodies.

Altogether, our results illustrate the power of U-ExM to decipher the molecular mechanisms behind LCA5, as well as to probe the impact at subcellar level of gene augmentation as a therapeutic strategy (summarized in Figure 7). In particular, our work unveiled the function of the bulge region defined by the LCA5 protein, whose depletion leads to abnormal axonemal structures, defective IFT and rhodopsin trafficking and severely affected OS membrane disc formation. We further show that *LCA5* gene augmentation appears as a promising therapeutical strategy that needs further optimization using notably the novel U-ExM pipeline that we describe in this paper.

**Figure 7.**
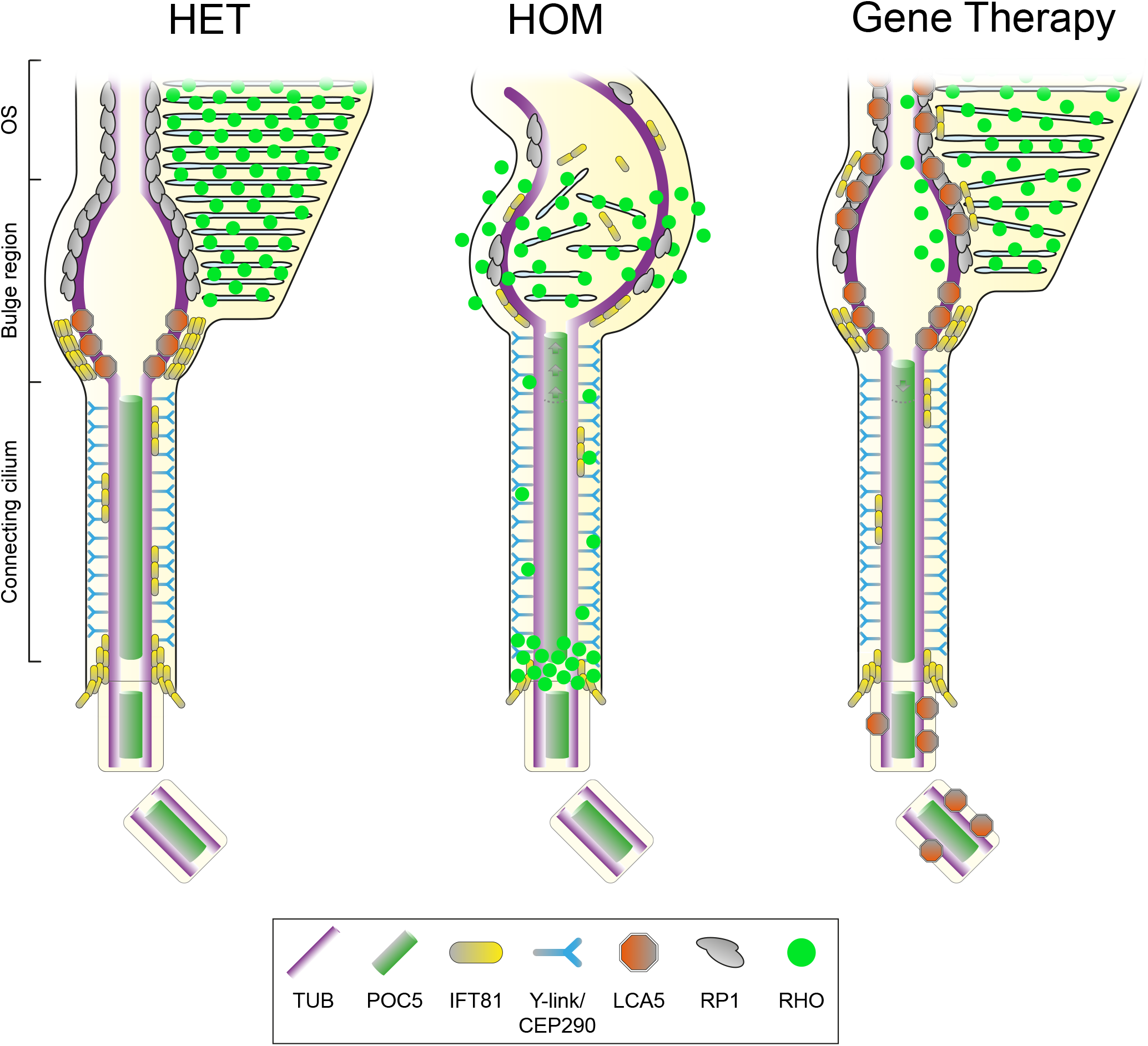
Schematic representation of *Lca5*^*+/gt*^ (HET), *Lca5*^*gt/gt*^ (HOM), and Gene Therapy treated photoreceptors. In unaffected HET photoreceptors (left), LCA5 (orange) localizes predominantly at the proximal part of the bulge region, between CEP290 (cyan, Y-links protein) and RP1 (grey, distal axonemal protein). IFT81 (yellow) localizes to the same bulge proximal region as LCA5, but also accumulates above the basal body and to a lesser extent along the CC. In HOM photoreceptors (middle), the CC is extended, as illustrated by an elongated CEP290 and POC5 (green) signal. Furthermore, bulge formation and distal axoneme organization is disrupted, leading to rhodopsin (RHO, bright green) mis-trafficking, with accumulation above the basal body, localization along the CC, and inside vesicle-like structures. Moreover, LCA5 loss leads to decreased levels of IFT81 and RP1 at the bulge region and more dispersed localization along the distal axoneme. AAV-*LCA5* gene augmentation therapy (right) partially restores bulge formation, CC and distal axoneme organization, as well as RP1 and rhodopsin localization. IFT81 localization is restored to a lesser extent, possibly explained by the ectopic LCA5 expression along the distal axoneme.

## Discussion

The sensory cilium of the photoreceptor cell is one of the longest cilia in the human body, notably due to the stacking of hundreds of opsin-loaded membrane discs. This causes a massive elongation of its tip, the outer segment, which is required for efficient photoreception and transduction. The importance of this unique sensory organelle is highlighted by the number of retinal diseases associated with mutations in genes coding for OS structure or function (37). The difficulty to characterize molecular and structural mechanisms involved in photoreceptor degeneration in such diseases mostly comes from the lack of resolution of current imaging tools. To overcome this problem, we recently adapted Ultrastructure Expansion Microscopy (U-ExM) for use in retinal imaging, enabling assessment of the photoreceptor cilium on regular fluorescence microscopes at unprecedented resolution. By this technique, we recently provided the first molecular mapping of the OS connecting cilium, that helped to understand molecular and structural mechanisms associated with a subtype of RP (13).

Here, we continued the molecular characterization of the outer segment focusing on the bulge region that directly extends from the connecting cilium (CC). We first describe that LCA5 decorates microtubule doublets (MTDs) externally on the proximal part of the bulge region. We also show that RP1, an LCA5 interactor that we revealed by affinity proteomics, localizes immediately distal to LCA5, extending its presence towards the axoneme distal end. Interestingly, we revealed that the bulge region corresponds to the location where rhodopsin-enriched membrane discs are formed, corroborating recent results suggesting that this region plays a role in the formation of membrane discs via an actin-dependent process involving PCARE and WASF3 proteins (15). Therefore, the bulge region appears as a strategic hub of the OS axoneme, explaining why mutations in genes encoding bulge proteins, such as LCA5, could have dramatic consequences on photoreceptor development and/or maintenance (1, 7). We previously demonstrated that this region exhibits a typical enlargement of the axonemal MTDs, whose function is yet to be determined. One could imagine that given the intense turnover of membrane discs, the bulge could act as a reservoir to maintain the constant amount of proteins needed to form membrane discs. In line with this, we also showed that the IFT-B machinery accumulates at the level of the bulge region, exactly where LCA5 is located, corroborating the interaction of these two modules at the bulge (7, 38, 39). One of the roles of LCA5 at the bulge could be to prevent IFT cargos to reach the more distal part of the axoneme, thus concentrating outer segment building blocks brought by IFT at the location where membrane discs form. Finally, given the microtubule-associated feature of LCA5, we also speculate that it could play a role on the structural maintenance of MTDs at the level of the bulge region.

Consistent with this model, we showed that LCA5 loss leads to rapid and drastic disorganization of the OS axoneme above the CC from P14 onwards, accompanied by loss of the bulge region. Surprisingly, CC and centriole structure remain unaffected, but CC length was significantly increased when the bulge was absent, suggesting that LCA5 could act as a ruler to dictate inner scaffold and CEP290 length. The loss of the bulge structure was accompanied by the loss of bulge-associated proteins, RP1 and IFT81, showing that their localization is dependent on LCA5. The concomitant loss of IFT81 at the level of the cilium entry could then be the consequence of a lack of protein recycling by IFT when the bulge is lost. In line with this, we showed that IFT81 signal at the bulge in *Lca5*^gt/gt^ (HOM) is progressively spread towards the distal axoneme before being lost, corroborating the role of LCA5 to stop IFT trains at the level of the bulge, allowing them to return to the cell body. The consequent outer segment collapse could result from the combination of the loss of cohesion between MTDs in the absence of LCA5, RP1 loss, that has been described to be required for proper OS disc orientation (30, 35), and IFT defects, possibly preventing the trafficking of OS building blocks.

Our study also explored, for the first time and with unprecedented resolution, the subcellular outcome of gene therapy using U-ExM. Therefore, our study provides additional critical information in sub-cellular transgene expression that is lacking in current pre-clinical gene therapy studies, making our approach a precious tool that could help to further optimize the efficacy of gene augmentation therapy.

In this study, we showed that AAV-*LCA5* injection in HOM retina leads to localization of LCA5 inside photoreceptor cells at the expected bulge region, similar to endogenous LCA5, but also decorating the entire distal axoneme and the basal body. We argue that this is due to the overexpression of *LCA5*, which in AAV-*LCA5* is under control of a strong promoter (CMV) (23). Whether the presence of LCA5 beyond the bulge region and throughout the axoneme could have deleterious effects on outer segment function remains an open question, but it may not be completely neutral. Importantly, affinity proteomics data from AAV-*LCA5* injected retina revealed enrichment of several centrosomal proteins, that we suggested to be associated to LCA5 centriolar localization upon ectopic overexpression. Whether these potential interactors could have a role on the LCA5-associated phenotype remains to be determined. However, we showed that exogenous LCA5 expression in photoreceptors partially rescues axoneme defects, RP1 and IFT protein localization, CC length, and is associated with improved photoreceptor survival. The relatively modest level of rescue could be explained by several arguments. First, the phenotype observed in HOM retina appears very early, between P10 and P14 onwards, whereas it has been described that exogenous LCA5 is expressed 15 days after intravitreal delivery, performed at P2. Second, intravitreal injection of AAV-*LCA5* leads to heterogeneous overexpression of LCA5 in virtually all retina layers, possibly leading to off-target or deleterious effects. Finally, LCA5 being a microtubule-binding protein, an important local cellular overexpression can impair microtubule network and lead to cell death. Altogether, these results show the importance of a precise and controlled spatiotemporal expression of LCA5 to improve gene therapy efficacy for LCA, that already showed interesting preliminary functional results (23, 24). With U-ExM, we now have the spatial resolution to accurately monitor rescue or induction of subcellular phenotypes, such as the axoneme reformation, or recruitment of interactors. U-ExM represents a powerful tool to assess and improve gene therapy efficacy in many cellular or tissue contexts, which is a crucial step to design preclinical studies before that a clinical trial can be initiated.

## Material and methods

### Mouse model

*Lca5*^*gt/gt*^ (HOM) and *Lca5*^*+/gt*^ (HET) (7) were handled in accordance with the statement of the “Animals in Research Committee” of the Association for Research in Vision and Ophthalmology, and experiments were approved by the Animal Ethics Committee (DEC) of the Radboud University Medical Center (AVD10300 2016 758; RU-DEC-2016-0050). The mice were maintained at RT with a 12-hour light/12-hour dark cycle with light on at 7:00 AM and were fed *ad libitum*. For affinity proteomics, mice were sacrificed at postnatal day ∼P28 of age and retinas were harvested. For ultrastructure expansion microscopy (U-ExM), mice were sacrificed at postnatal (P) day P10, P14, P18, P22, and P28.

### Intravitreal (IVT) injections

IVT injections were performed similarly as described before (40). Briefly, postnatal day (P)2 HOM and HET mouse pups of either sex were anesthetized by hypothermia. Eyelids were opened using a 30- gauge needle. An incision was made into the sclera at the nasal part of the retina. For affinity proteomics, 0.6 μl of AAV7m8.CbA.hopt-*LCA5* (9.87E+9 vg/μl) (23), containing a C-terminal 3xFLAG-tag, was injected in one eye using a Hamilton syringe, while 0.6 μl of AAV7m8.CbA.*eGFP* (control vector; 9.98E+9 vg/μl) (23) was injected in the other eye as an internal control. For U-ExM, 0.6 μl of AAV7m8.CbA.hopt-*LCA5* (9.87E+9 vg/μl) (23), containing a C-terminal 3xFLAG-tag, was injected in one eye using a Hamilton syringe, while the other eye was left untreated to not interfere with the morphological development.

For affinity proteomics, a total of 15 mice for each genotype (HOM and HET) were injected per biological replicate (n=5). For U-ExM, a total of 15 mice for each genotype (HOM and HET) were injected, using three mice of each genotype per timepoint (P10, P14, P18, P22, and P28).

### Affinity proteomics on mouse retina

Four weeks after IVT injections, retinas were harvested by making a cut in the eye with removing the lens, followed by carefully squeezing out the retina with a forceps. Harvested retinas were pooled per group (HOM + AAV-*LCA5*; HOM + AAV-*eGFP*; HET + AAV-*LCA5*; HET + AAV-*eGFP*). Retinas were lysed in lysis buffer containing 0.5% Nonidet-P40, protease inhibitor cocktail (Roche), and phosphatase inhibitor cocktails II and III (Sigma-Aldrich) in TBS (30 mM Tris-HCl, pH 7.4, and 150 mM NaCl) for 30 minutes at 4 °C, followed by high-speed shaking (30Hz) using Tissuelyser II (Qiagen) and sonication (>20 kHz; 10 cycles – 30 sec ON/30 sec OFF) at 4 °C. Lysates were cleared by centrifugation at 10,000 xg for 15 minutes. Cleared lysates with equal amounts of proteins were transferred to anti-FlagM2 affinity gel (Sigma-Aldrich) and incubated for 1.5 hours at 4 °C. Subsequently, the affinity gel with bound protein complexes were washed three times with wash buffer (TBS containing 0.1% NP40 and phosphatase inhibitor cocktails II and III), followed by two times with 1x TBS. On-bead digestion was performed for 30 minutes at 27 °C and 800 rpm, in Trypsin digestion buffer (2 mM Urea, 50 mM Tris-HCl (pH 7.5), 5 μg/sample Trypsin (Serva, 37283)). The supernatant was collected and the beads were washed once with urea/DTT buffer (2 mM urea, 50 mM Tris-HCl (pH 7.5), 1 mM DTT). The on-bead digestion and wash were pooled per sample and incubated overnight (ON) at room temperature (RT). The next day, samples were snap frozen and stored at -80 °C until mass spectrometry analysis.

### Mass spectrometry analysis, protein quantification and statistics

The trypsinized protein samples were subjected to Iodacetamide (IAA, 5 mg/ml, Merck, 8.04744.0025) and StageTip purified (Thermo Scientific), followed by mass spectrometry analysis on an Orbitrap Fusion Tribrid mass spectrometer (Thermo Fisher) as described earlier (39). Label-free quantification (LFQ) was performed, using Maxquant (v.1.6.1.0) and identified proteins were analyzed using Perseus (v1.6.2.3) (41, 42). This procedure was performed twice per sample and combined in one dataset using the mean values. All data were filtered for potential contaminants, peptides only identified by site, and reverse database identifications. The proteins were filtered to be present in ≥ 3 of the 5 replicates. For AAV-*LCA5* versus AAV-*eGFP* (control) injected retinas, a one-sided two-sample test with was performed (p<0.05). Furthermore, a significance A outlier test with permutation-based FDR correction was performed (FDR<0.05). Proteins were considered significantly enriched when passing both the two-sample test (p<0.05) and the significance A outlier test (FDR<0.05). Datasets from HOM and HET mice were analyzed separately.

### Ultrastructure Expansion Microscopy (U-ExM) on mouse retina

Retinas were prepared for U-ExM, as earlier described (13). In short, eyes of P10, P14, P18, P22, and P28 mice were enucleated and fixed for 15 minutes at RT in 4% Paraformaldehyde (PFA)/PBS. Subsequently, the cornea and lens were removed with micro-scissors followed by carefully removing the retina. The retinas were incised to flatten it and placed inside a 10 mm microwell of a 35 mm petri dish (P35G-1.5-10-C, MatTek) for U-ExM processing.

The expansion procedure is an adaptation of the original U-ExM protocol that we have optimized for eye tissue samples as previously described (13, 43). Briefly, retinas were incubated ON in 100 μl of 2% acrylamide (AA; A4058, Sigma-Aldrich) + 1.4% formaldehyde (FA; F8775, Sigma-Aldrich) at 37 °C. The next day, retinas were incubated in 35 μl monomer solution composed of 25 μl of sodium acrylate (stock solution at 38% (w/w) diluted with nuclease-free water, 408220, Sigma-Aldrich), 12.5 μl of AA, 2.5 μl of N,N’-methylenebisacrylamide (BIS, 2%, M1533, Sigma-Aldrich), and 5 μl of 10× PBS for 90 minutes at RT. Subsequently, the monomer solution was removed and retinas were incubated in 90 μl of monomer solution with the addition of ammonium persulfate (APS, 17874, Thermo Fisher Scientific) and tetramethylethylenediamine (TEMED, 17919, Thermo Fisher Scientific) as a final concentration of 0.5% for 45 minutes at 4 °C followed by 3 hours incubation at 37 °C. A 24-mm coverslip was added on top to close the chamber. Next, the coverslip was removed and 1 ml of denaturation buffer (200 mM SDS, 200 mM NaCl, 50 mM Tris Base in water (pH 9) was added into the MatTek dish for 15 minutes at RT with shaking to detach the gel from the dish. Afterwards, the gel was incubated in denaturation buffer for 1 hour at 95 °C followed by ON at RT. The next day, the gel was sliced around the retina that is still visible at this step, and expanded in three consecutive ddH_2_O baths. Next, the gel was manually sliced with a razorblade to obtain approximately 0.5 mm thick transversal sections of the retina to enable processing for immunostaining.

For immunostaining, primary antibodies (Supplemental Table 2) were incubated ON at 4 °C or for 3 hours at 37 °C for anti-RP1. Image acquisition was performed on an inverted Leica Thunder DMi8 microscope using a 20x (0.40 NA) or 63x (1.4 NA) oil objective with Thunder SVCC (small volume computational clearing) mode at max resolution, adaptive as “Strategy” and water as “Mounting medium” to generate de-convolved images. Z-stacks were acquired with 0.21 μm z-intervals and an x, y pixel size of 100 nm.

### Human cell culture and expansion

U2OS cells were grown at 37 °C with 5% CO_2_ in DMEM supplemented with GlutaMAX (Life Technology), 10% fetal calf serum (Brunschwig), penicillin and streptomycin (100 μg/ml). For U-ExM, U2OS cells were plated onto coverslips in a 6-well plate at 300,000 cells/well and processed 24 hours later. Briefly, coverslips were incubated for 3 hours in 100 μl of 2% acrylamide (AA; A4058, Sigma-Aldrich) + 1.4% formaldehyde (FA; F8775, Sigma-Aldrich) at 37 °C. Then, coverslips are incubated in monomer solution (10% AA, 19% SA, 0.1% BIS in 1x PBS) containing 0.5% TEMED and APS for 5 minutes on ice followed by 1 hour at 37 °C. Next, the gel is detached from the coverslip by adding 1 ml of denaturation buffer (200 mM SDS, 200 mM NaCl, 50 mM Tris Base in water (pH 9) for 15 minutes under shaking into a well of a 6-well plate. The gel is then transferred to a 1.5 ml tube and incubated in denaturation buffer for 1.5 hour at 95 °C. Gels were washed from the denaturation buffer twice in ddH2O. Then, gels were shrunk in 1x PBS and stained for 3 hours at 37 °C in 1x PBS-BSA 2% for both primary (Supplemental Table 2) and secondary antibodies (1:400), followed each by 3 washes for 5 minutes with 1x PBS-Tween 0.1%. Finally, gels were next re-expanded in ddH_2_O and imaged.

### Quantifications

#### Expansion factor

The expansion factor was calculated in a semi-automated way by measuring the full width at half maximum (FWHM) of photoreceptor mother centriole proximal tubulin signal using PickCentrioleDim plugin, as described before (13). A total of 100 photoreceptor mother centrioles, divided over both genotypes, all timepoints, and all three replicates, were subjected to FWHM measurements and compared to a pre-assessed value of U2OS centriole width (mean = 231.3 nm +/− 15.6 nm) (13). Calculating the ratio between measured FWHM and known centriole width resulted in an expansion factor of 4.39 (Supplemental Table 3). This expansion factor was then used for every quantification. Scale bars shown in all figures are corrected for the expansion factor.

#### Protein signal length and position

Protein signal lengths or position compared to basal body proximal end (depicted with tubulin) were measured using a segmented line drawn by hand (FIJI) to fit with photoreceptor curvature, and corrected with the expansion factor. Only photoreceptors where both protein signal (POC5, CEP290, LCA5, RP1, or IFT81) and basal body proximal end (tubulin) were clearly visible were selected for measurement.

#### Tubulin spread

Tubulin spread was assessed at P10, P14, P18, and P22 in non-injected retinas and at P18 in injected retinas in a semi-automated way by measuring FWHM of tubulin signal with PickCentrioleDim plugin (13) on 3 different locations of the photoreceptor corresponding to 500 nm proximally to the end of the connecting cilium (CC) POC5 staining (−500), at the level of the end of the CC POC5 staining (0), or 500 nm distally to the end of the CC POC5 staining (+500). For injected photoreceptors, the proximal extremity of the expressed LCA5 signal was used to set the (0) location. Each measurement was subsequently corrected for the expansion factor.

#### Fluorescence intensity

Fluorescence intensity measurements of RP1, Rhodopsin, and IFT81 were performed on maximal projections using FIJI (44) on deconvoluted images. The same rectangle region of interest (ROI) drawn by hand was used to measure the mean gray value of the protein signal and their corresponding background. Fluorescence intensity was finally calculated by dividing the mean gray value of the fluorescence signal by the mean gray value of the background (normalized mean gray value). For RP1, measurements were performed on the bulge region, defined by tubulin. For Rhodopsin, measurements were performed on the outer nuclear layer (ONL) and on the OS layer on 20x images. For each image, 3 different ROI are measured for ONL and OS layers and fluorescence intensity was calculated by dividing average ONL mean gray value by OS average mean gray value. For IFT81, measurements were performed on the region just above the basal body (BB) and on the bulge region, defined by tubulin.

#### Axonemal defects

Axonemal defects were categorized as follows. Photoreceptors showing axoneme bending over 180° were classified as “Bent/Curled” whereas photoreceptors exhibiting microtubule spreading or even loss of distal axoneme were classified as “Open/Broken”.

#### ONL thickness

ONL thickness was measured manually using tubulin and/or DAPI staining on two different 20x images per replicate. Three different measurements were performed per image to avoid bias due to retina dissection. Each measurement was subsequently corrected for the expansion factor.

#### Exogenous LCA5 expression in injected retinas

The percentage of photoreceptors expressing LCA5 was quantified manually from 63x images, independent of the level of expression.

#### Statistical analyses

The comparison of 2 groups was performed using non-parametric Mann–Whitney test, if normality was not granted, because rejected by Pearson test. Every measurement was performed on 3 different animals. Data are all represented as a scatter dot plot with centerline as mean, except for percentages quantifications, which are represented as histogram bars. For tubulin width measurements, the variances of each condition (HET, HOM, HOM + Therapy) and at every location (−500, 0, 500) were compared using an F-Test. The graphs with error bars indicate SD (+/−) and the significance level is denoted as usual (non-significant (ns) p>0.05, * p < 0.05, ** p < 0.01, *** p < 0.001, ****p < 0.0001). All the statistical analyses were performed using Graphpad Prism9. Every mean, standard deviation, test, and the number of animals used for comparison are referenced in Supplemental Table 3. When possible, a minimum of 10 measurements has been performed per animal. The data underlying the graphs shown in all the figures are included in Supplemental Table 3.

## Author contributions

S.F. and O.M. participated in experimental design, performed research, collected, analyzed, and interpreted data, performed statistical analysis, and drafted and revised the manuscript. S.F. and O.M. share the first-author position, with S.F. as first-first since S.F. and R.R. initiated the project. K.J. processed samples after affinity purification to perform mass spectrometry analysis. A.G. assisted in the intravitreal injections, co-supervised the project, and revised the manuscript. M.U. was responsible for the funding acquisition and supervision of the mass spectrometry analysis. R.W.J.C. supervised the project, interpreted data and revised the manuscript. K.B. supervised the mass spectrometry analysis, analyzed and interpreted data, and revised the manuscript. P.G., V.H., and R.R. were responsible for funding acquisition and supervision of the project, interpreted data, and revised the manuscript. P.G., V.H., and R.R. share the last-author position, with R.R. as last author since R.R. initiated the project. The position of P.G. and V.H. was determined alphabetically.

## Acknowledgments

We thank Prof. Jean Bennett for providing the AAV vectors; Prof. Helen May-Simera for training SF to perform the intravitreal injections; and Qin Liu and Rossano Butcher for providing the RP1 antibody.

This research was funded by the Landelijke Stichting voor Blinden en Slechtzienden, Stichting Retina Nederland Fonds, Stichting Beheer Het Schild, Stichting Blinden-Penning, and Stichting Steunfonds Uitzicht via Uitzicht 2016/2017-22, together with the Gelderse Blindenstichting, Rotterdamse Stichting Blindenbelangen, Stichting tot Verbetering van het Lot der Blinden ‘Het Lot’, the Stichting voor gehandicapte kinderen Dowilvo, the Netherlands Organisation for Health Research and Development (ZonMW, #91216051), and the Foundation Fighting Blindness, USA (PPA-0717-0719-RAD and BR-CMM-0720-0789-RAD) to R.R. This work is also supported by the Swiss National Foundation (SNSF) 310030_205087 and the Pro Visu Foundation attributed to P.G. and V.H. M.U. and K.B. are supported by the Tistou & Charlotte Kerstan Stiftung.

