## Supplementary material for "Probing the sub-cellular mechanisms of LCA5-Leber Congenital Amaurosis and associated gene therapy with expansion microscopy": Supplemental_Materials.pdf

**A**

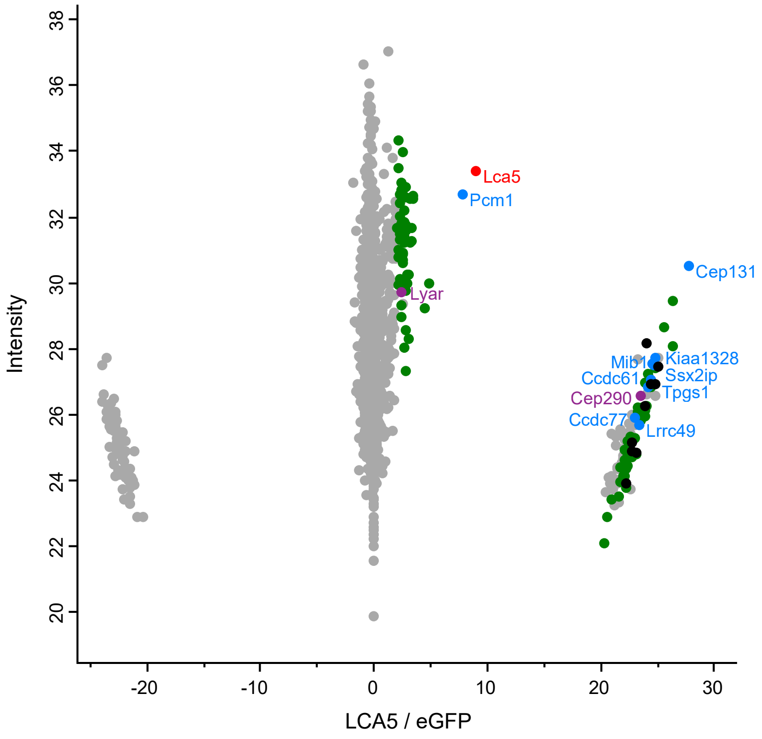


**B**

| Category | Protein names |
|---|---|
| Photoreceptor proteins | Cep290 Lyar |
| Centriolar satellite/ Centrosomal proteins | Ccdc61 Ccdc77 Cep131 Kiaa1328 Lrrc49 Mib1 Pcm1 Ssx2ip Tpgs1 |
| Ribonucleo-proteins | [Supplemental Table 1] |
| Miscellaneous | [Supplemental Table 1] |

**Supplemental Figure 1. Identification and clustering of potential LCA5 interactors.** (**A**) Scatterplot showing enriched proteins comparing AAV-*LCA5* injected retinas to AAV-*eGFP* (control) injected retinas in unaffected *Lca5*$^{+/gt}$ (HET) mice. The bait protein (LCA5) is shown in red. Significantly enriched proteins are categorized into different groups based on their function, including photoreceptor-associated proteins (purple), centriolar satellite/ centrosomal proteins (blue), ribonucleoproteins (green), and miscellaneous (black). X-axis represents log2 ratio between AAV-*LCA5* and AAV-*eGFP* (control) injected retinas. Y-axis represents the intensity score, indicating the relative amount of proteins in the dataset. (**B**) Table showing the significant proteins categorized in different groups, based on their function. Original data is listed in Supplemental Table 1.

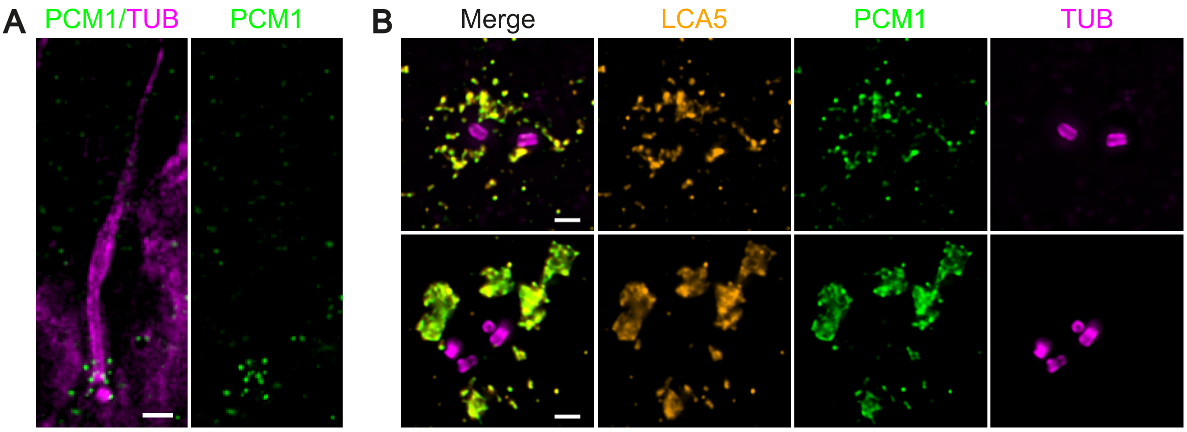


**Supplemental Figure 2. Localization of PCM1 in photoreceptor cells and human U2OS cultured cells.** (**A**) Widefield (63x) images of expanded adult photoreceptor stained for tubulin (magenta) and PCM1 (green). Scale bar: 500 nm. (**B**) Representative widefield (63x) images of expanded U2OS centrioles stained with tubulin (magenta), LCA5 (orange), and PCM1 (green). Scale bars: 500 nm.

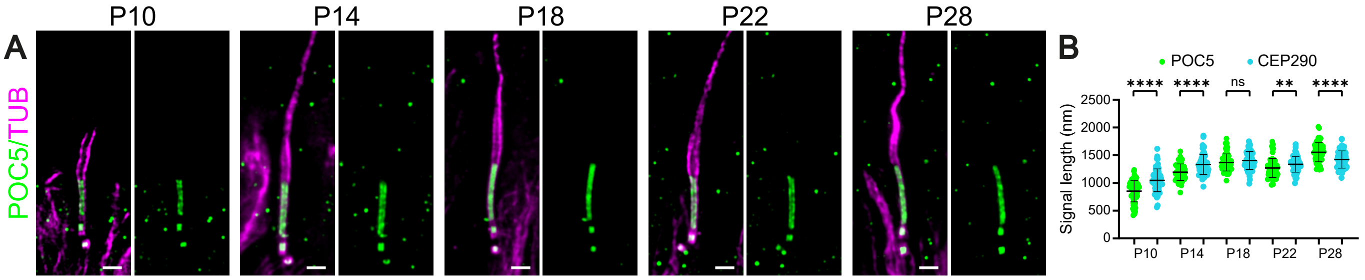


**Supplemental Figure 3. Localization of POC5 in developing photoreceptors.** (**A**) Widefield (63x) images of expanded photoreceptors stained for tubulin (magenta) and POC5 (green) from P10 to P28 in *Lca5$^{+/gt}$* (HET) mice. Scale bars: 500 nm. (**B**) Comparison of POC5 (green) and CEP290 (cyan) signal length from P10 to P28 in HET mice. Note that these measurements correspond to the data in Figure 3, G and H. Three animals per timepoint.

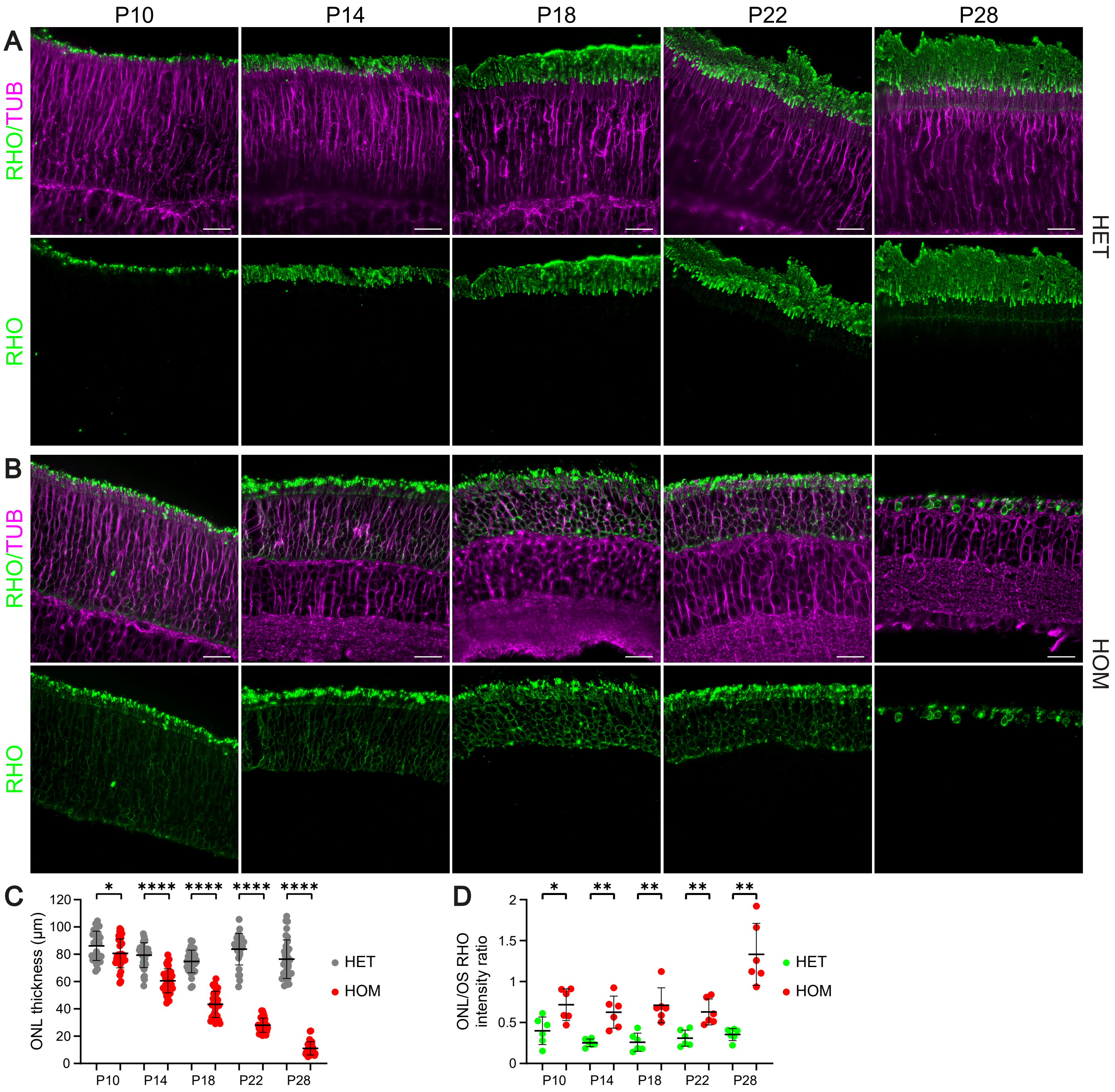


**Supplemental Figure 4. Development of retinal layers in *Lca5*$^{+/gt}$ (HET) and *Lca5*$^{gt/gt}$ (HOM) mice.** (**A-B**) Low magnification (20x) widefield images of expanded retinas in HET mice (A) and HOM mice (B) showing rod OS development from P10 to P28 by staining with Rhodopsin (green) and tubulin (magenta). Scale bars: 20 µm. (**C-D**) Impact of LCA5 loss on outer nuclear layer (ONL) thickness (C) or ONL/OS Rhodopsin intensity ratio (D) from P10 to P28. Note that the HET and HOM measurements of P18, P22, and P28 correspond to the data in Figure 6, D and E. Three animals per timepoint.

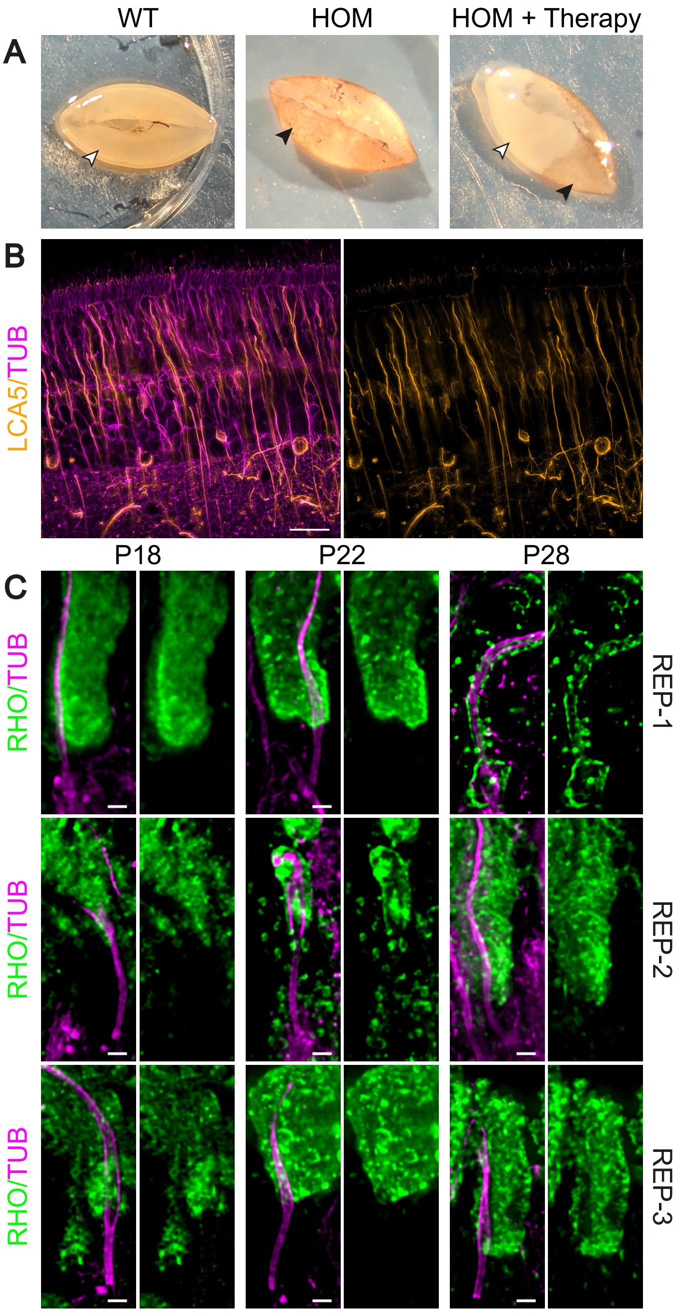


**Supplemental Figure 5. Global effect of AAV-*LCA5* gene augmentation therapy.** (**A**) Representative pictures of dissected retinas from wild-type (WT), *Lca5*$^{gt/gt}$ (HOM), and AAV-*LCA5* gene therapy treated HOM mice. Lack of pigmentation in WT and HOM + therapy retinas is indicated by open arrowheads and pigmentation in HOM and HOM + therapy retinas is indicated by closed arrowheads. (**B**) Low magnification (20x) widefield images of expanded P28 retina, treated with AAV-*LCA5* gene therapy and stained for LCA5 (orange) and tubulin (magenta). Scale bar: 20 µm. (**C**) Representative widefield (63x) images of expanded photoreceptors of each replicate (REP1-3) stained for tubulin (magenta) and Rhodopsin (green) from P18 to P28 in AAV-*LCA5* gene therapy treated HOM mice. Of note: Each replicate represents a different mouse. Scale bars: 500 nm.

**Supplemental Table 1. Protein list of potential LCA5 interactors.** (**A**) Table showing significantly enriched proteins in AAV-*LCA5* injected retinas compared to AAV-*eGFP* (control) injected retinas in *Lca5*$^{gt/gt}$ (HOM) mice (**Tab 1**) and unaffected *Lca5*$^{+/gt}$ (HET) mice (**Tab 2**). The bait protein (LCA5) is shown in red. Significantly enriched proteins are categorized into different groups based on their function, including photoreceptor-associated proteins (purple), centriolar satellite/centrosomal proteins (blue), ribonucleoproteins (green), and miscellaneous (black).

**Supplemental Table 2. Antibody list.**

**Supplemental Table 3. Descriptive statistics for all measurements.**
